# Predictors of sensorimotor adaption: insights from over 100,000 reaches

**DOI:** 10.1101/2023.01.18.524634

**Authors:** Jonathan S. Tsay, Hrach Asmerian, Laura T. Germine, Jeremy Wilmer, Richard B. Ivry, Ken Nakayama

## Abstract

Sensorimotor adaptation is essential for keeping our movements well-calibrated in response to changes in the body and environment. For over a century, we have studied sensorimotor adaptation in highly controlled laboratory settings that typically involve small sample sizes. While this approach has proven useful to characterize different learning processes, laboratory studies are typically very underpowered to generate data suited for exploring the myriad of factors that may modulate motor performance. Here, using a citizen science website (testmybrain.org), we collected over 2000 sessions on a visuomotor rotation task. This unique dataset has allowed us to replicate classic motor findings, reconcile controversial findings in the learning and memory literature, and discover novel constraints underlying dissociable implicit and explicit learning processes supporting sensorimotor adaptation. Taken together, this study suggests that large-scale motor learning studies hold enormous potential to advance sensorimotor neuroscience.

## Introduction

Sensorimotor adaptation keeps our movements well-calibrated in response to changes in the body and environment. For example, sensorimotor adaptation can help a tired basketball player compensate for their muscle fatigue and accelerate recovery in patients undergoing neurorehabilitation (1–3).

The study of sensorimotor adaptation traces back to the early days of experimental psychology (4,5): For example, in 1897, George Stratton published his classic self-experiment, describing the behavioral and psychological changes he experienced when wearing mirror-inverting glasses for eight consecutive days. In the 21^st^ century, these questions are typically addressed by using environments and virtual reality systems that allow the experimenter to perturb the movement feedback (6–9). For example, a visuomotor perturbation can be introduced by rotating the position of the cursor from the actual hand position. The mismatch between the expected and actual position of the visual feedback elicits adaptation, that is, movements in the opposite direction of the rotation that reduce and eventually nullify the visuomotor error. If the perturbation is small, this change in hand angle emerges gradually and occurs outside the participant’s awareness, a phenomenon known as implicit recalibration (10). If the perturbation is large, the adaptive response may be accompanied by more explicit adjustments in aiming (11–16).

Studies of sensorimotor adaptation are typically conducted with specially designed apparatuses in controlled laboratory settings. This approach has been extremely successful in revealing critical spatial (17–20) and temporal (21–25) constraints on adaptation, as well as examining the contributions of different neural systems to this form of learning (1,2,12,26–32). However, in-person research typically involves small, homogenous samples (33), making certain research questions impractical and difficult to answer (e.g., exploring how different demographic factors modulate motor behavior). Moreover, analyses that identify potential causal underpinnings using small datasets are susceptible to overfitting, which increases the likelihood that findings fail to generalize outside the experimental context and to different populations (34).

To address these issues, we designed a web-based visuomotor rotation task (35,36) and collected more than 2000 sessions of data through a citizen science website (www.testmybrain.org) (37–40). Leveraging this unique dataset (41), we built a cross-validated, predictive model to identify factors that support successful motor adaptation. The results from this study not only replicate classic findings in the literature but also highlight novel, widely generalizable constraints underlying sensorimotor learning.

## Results and Discussion

### The viability of studying sensorimotor adaptation outside the traditional laboratory

We collected 2,121 sessions of data through the testmybrain.org website. The data included results from a demographic survey and behavior from a web-based visuomotor rotation task. Participants completed different numbers of sessions and we employed a few variants of the task (Methods: Table 1-2).

**Table 1.** 2,121 total experimental sessions broken down by session number and target version. 44 sessions were not identified due to an error in our survey configuration.

| Target # / Session # | 1<sup>st</sup>-session | 2<sup>nd</sup>-session | 3<sup>rd</sup>-session | 4<sup>th</sup>-session | Not identified | Total |
| --- | --- | --- | --- | --- | --- | --- |
| 1-target | 1,747 | 80 | 13 | 23 | 41 | 1,904 |
| 2-target | 181 | 7 | 2 | 2 | 3 | 195 |
| 4-target | 10 | 1 | 0 | 1 | 0 | 12 |
| 8-target | 10 | 0 | 0 | 0 | 0 | 10 |
| Total | 1948 | 88 | 59 | 25 | 44 | 2,121 |

**Table 2:** Summary of demographic and task features. The mean age (min – max) is provided. The mean and SDs are provided for self-reported Likert ratings of clumsiness (“I am clumsy”) and self-reported Likert ratings of overall experience completing the experiment (“I enjoyed the experiment”). A rating of 1 and 5 signified that the participant strongly disagreed or strongly agreed with the statement, respectively. Median splits for baseline variability (°), reaction time (ms), movement time (ms), and search time (ms) are provided.

| Total n = 1, 747 (first experimental session; one-target version) |  |  |  |  |  |
| --- | --- | --- | --- | --- | --- |
| **Age** |  | 26.3 (9 – 96) | **Sex** | Female<br>Male<br>Other | 855<br>839<br>53 |
| **Rating of clumsiness** |  | 3.0 (1.1) | **Vision intact** | Yes<br>No | 1584<br>163 |
| **Rating of enjoyment** |  | 3.3 (1.1) | **Screen size** | Height<br>Width | 787.3 (160.7)<br>1525.8 (319.1) |
| **Hours of computer use** |  | 6.9 (3.2) | **Browser** | Chrome | 1375 |
| **Hours of sleep per night** |  | 7.1 (1.4) |  | Firefox<br>Edge<br>Opera<br>Safari<br>Not detected | 142<br>3<br>42<br>177<br>8 |
| **Handedness** | Right | 1547 | **Target location** | Cardinal | 835 |
|  | Left<br>Ambidextrous | 150<br>50 |  | Diagonal | 912 |
| **Device** | Mouse | 764 | **Undergraduate Major** | STEM | 695 |
|  | Trackpad<br>Trackball<br>Other | 942<br>18<br>23 |  | Psychology<br>Social Science<br>Business<br>Arts/Humanities<br>Other | 208<br>108<br>162<br>201<br>373 |
| **Racial origin** | Multi-racial | 79 | **Highest level of education** | Primary | 21 |
|  | White<br>Asian<br>Latinx<br>African American<br>Native American<br>Pacific Islander<br>Other<br>Rather not say | 833<br>431<br>116<br>55<br>22<br>5<br>93<br>113 |  | Middle<br>Secondary<br>Some college<br>Technical school<br>Bachelor<br>Graduate<br>Other<br>Rather not say | 286<br>358<br>388<br>54<br>313<br>262<br>11<br>54 |
| **Baseline variability** | High | 5.4 (2.1) | **Movement Time** | Fast | 97.4 (27.6) |
|  | Low | 2.8 (0.7) |  | Slow | 267.7 (130.8) |
| **Reaction Time** | Fast | 231.2 (37.0) | **Search Time** | Fast | 1384.5 (151.5) |
|  | Slow | 348.5 (59.8) |  | Slow | 1897.2 (273.4) |

We first focused on naïve participants who completed a task in which all reaches were to a single target (# of sessions = 1,747): After a familiarization block with veridical feedback, a 45° visuomotor rotation was imposed between the participant’s movement and visual cursor feedback (Figure 1a). To compensate for this rotation, participants exhibited significant changes in hand angle in the opposite direction of the rotation, gradually drawing the cursor closer to the target (Figure 1b). Individuals exhibited changes in hand angle during both early (Mean ± SEM: 22.3° ± 0.3°) and late phases of adaptation (Mean ± SEM: 34.5° ± 0.3°) (Figure 1d). When asked to forgo the use of any strategy-based change in behavior and reach directly to the target without visual feedback, participants exhibited robust aftereffects – a signature of implicit recalibration (Mean ± SEM: 12.6° ± 0.2°). Together, these data show a striking resemblance with those collected in the lab (35) (also see (42,43)).

**Figure 1.**
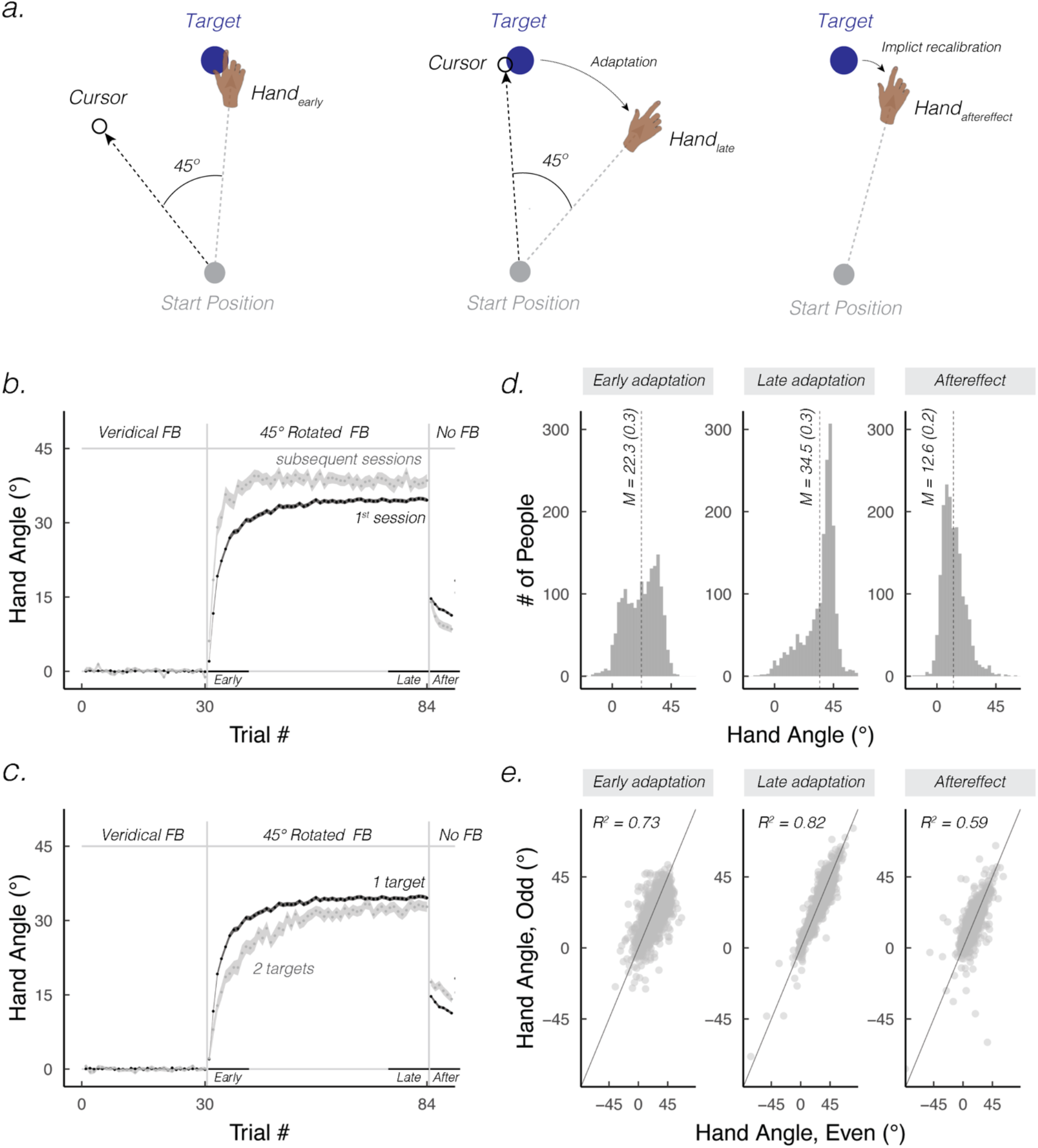
Web-based sensorimotor adaptation task and behavior. **(a)** Schematic of the sensorimotor adaptation task. The cursor feedback (white dot) was rotated 45° with respect to the movement direction of the hand. Participants were instructed to move such that the cursor would intersect the target (blue circle). Left, middle, and right panels display hand and cursor positions during the early, late, and aftereffect phases of learning, respectively. **(b)** Mean learning functions from naïve participants who completed the one-target version of the task for the first time (black; # of sessions = 1,747) vs. non-naïve participants completing the one-target version subsequent times (grey; # of sessions = 157). Shaded region represents SEM. **(c)** Mean time courses of hand angle for naïve participants who completed the one-target (black; n = 1,747) or two-target version (grey; n = 181) of the task. A hand angle of 0° denotes a movement directed to the target. **(d)** Distribution of participants’ mean hand angles during early, late, and aftereffect phases. **(e)** Split-half reliability correlating hand angles on even and odd trials across all three phases. Grey dots denote individual participants; grey lines represent the identity line.

There were substantial individual differences in hand angle across the three phases (Figure 1d). Given the limited time available for each participant, we compared odd and even trials as an assessment of reliability. The split-half reliability was moderate to high across all three phases (Figure 1e; Early: *R*^2^= 0.58, *ICC* = 0.72, *p* < 0.001; Late: *R*^2^ = 0.86, *ICC* = 0.93, *p* <0.001; Aftereffect: *R*^2^ = 0.59, *ICC* = 0.77, *p* <0.001). These results are especially noteworthy given that participants performed the task without any supervision.

We next examined how individual differences in hand angle correlated across the three phases (Figure 2). The change in hand angle during early and late phases of adaptation were highly correlated (Early vs Late: *R* = 0.59, *p* <0.001), whereas both measures were weakly correlated with that of the aftereffect phase (Early vs Aftereffect: *R*= − 0.05, *p* = 0.04; Late vs Aftereffect: *R*= −0.03, *p* = 0.25). These results are consistent with the classic idea that adaptation and aftereffect phases tap into dissociable learning processes, with the former including a substantial contribution from explicit strategic use (i.e., re-aiming) and the latter primarily reflective of implicit recalibration (44).

**Figure 2.**
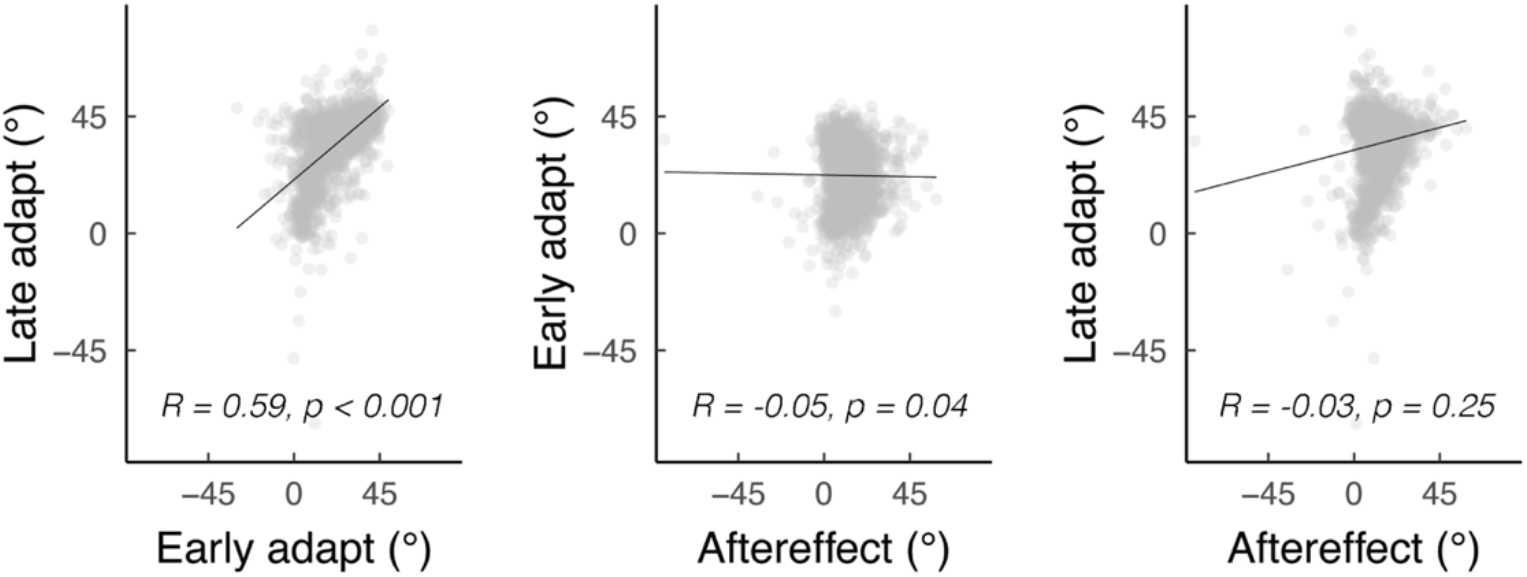
Cross-correlations between early adaptation, late adaptation, and aftereffect phases. *R* denotes Spearman’s correlation.

The data also replicate two classic effects in the sensorimotor adaptation literature. First, repeated exposure to the same, large visuomotor rotation has been shown to enhance the rate of adaptation but attenuate the size of the aftereffect (45–50). The former is attributed to the recall of a successful re-aiming strategy (49), whereas the mechanism for the latter remains an open question (45). To quantify these effects in our data, we compared the learning functions between participant’s first session (# of sessions = 1,747) and subsequent sessions (i.e., # of sessions = 157) (Table 1). There was a significant phase x session interaction (*F*(2, 5582) = 26.6, *p* <0.001) (Figure 1b): In subsequent sessions, an increase was observed for both early adaptation (*t*(5582) = 8.0, *p* <0.001) and late adaptation (*t*(5582) = 3.5, *p* <0.001). However, the aftereffect was reduced (*t*(5582) = 2.2, *p* = 0.02) (Figure 3a).

**Figure 3.**
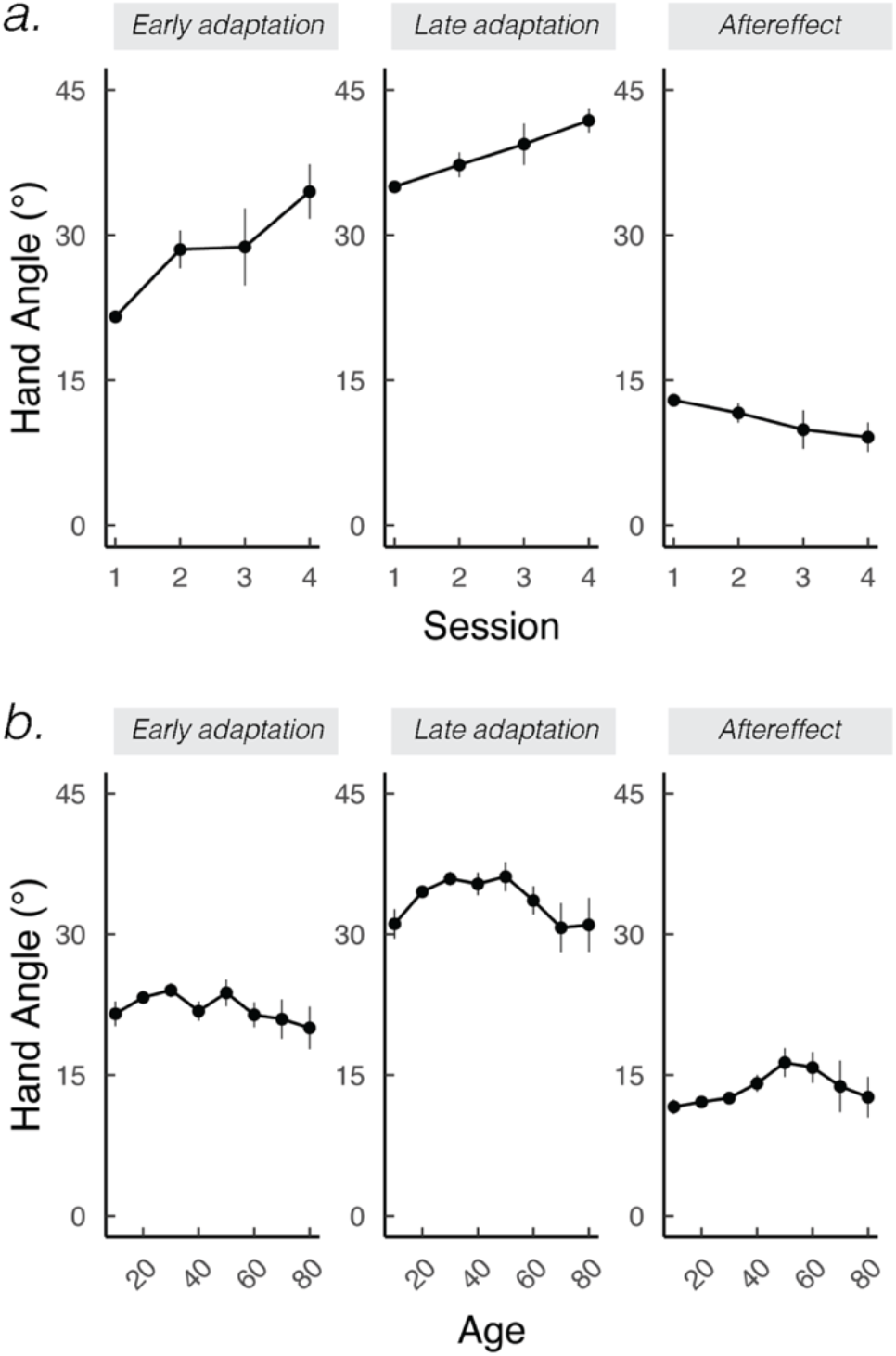
The effect of session and age on sensorimotor adaptation. **(a)** With exposure to the same visuomotor rotation across sessions (# of sessions = 1,863; Table 1), participants exhibit increased early and late adaptation across sessions but an attenuated aftereffect. **(b)** The inverted-U effect of age. For ease of visualization, participants’ age was binned based on increments of 10. Error bars denote SEM.

Second, the data replicate classic contextual interference effects in the learning and memory literature (Figure 1c) (25,51–55). Specifically, contextual interference is a phenomenon where learning is slower but better retained in a random practice schedule compared to a blocked practice schedule (25,56). The constant switching between contexts in random practice is thought to create more elaborate and better retained motor memories (51,54,57). To evaluate this phenomenon in our data, we compared the learning functions of naïve participants (i.e., 1^st^ session) who were tested on either a one-target (# of sessions = 1,747) or two-target (# of sessions = 181) version of our task (Figure 1c and see also Table 1). Note that the former is, by definition, blocked (i.e., there is only one target), whereas in the latter, the two targets are interleaved. There was a significant phase x version interaction (*F*(2, 5777) = 23.5, *p* <0.001): For the two target version, early adaptation (*t*(5777) = 6.0, *p* <0.001) and late adaptation (*t*(5777) = 2.1, *p* <0.001) were attenuated, but the aftereffect was larger (*t*(5777) = 3.6, *p* <0.001), reproducing the canonical signature of contextual interference.

These data also bear on a controversy in the motor learning literature concerning the effect of aging on sensorimotor learning. Several studies have reported no effect of age on motor adaptation (58–60). Others have found that aging impairs late adaptation (61–65), with this attenuation attributed to an age-related decline in strategy use. However, these previous studies have recruited modest sample sizes drawn from a limited age range (58,66). Leveraging our large dataset that spans a wide age range, we discovered a striking, inverted-U effect of age on all phases (quadratic AIC – linear AIC: early, Δ*AIC* = −5.2; late, Δ*AIC* = −17.2; aftereffects, Δ*AIC* = −1.1; all p < 0.05) (Figure 3b). Interestingly, late adaptation reached a peak at around 20 and dropped off at 60 years old; in contrast, the size of the aftereffect reached a peak around 50 and dropped off at 70 years old. The mixed results in the literature may be due to different studies sampling different points on the inverted-U curve (also see: (67)).

Taken together, the data indicate that our web-based visuomotor task yields data that are valid and reliable, replicate classic effects in the motor learning literature, and can elucidate unappreciated constraints on sensorimotor adaptation.

### Identifying predictors of explicit strategic re-aiming

By collecting a range of demographic and kinematic variables in a large sample, we are positioned to identify predictors of sensorimotor adaptation and forecast sensorimotor behavior that has not yet been observed (34). To this end, we adopted a machine learning approach that segregated our large sample of naïve participants who completed the one-target version on their first session (n = 1,747) into independent model development and testing subsets. Our cross-validated group lasso procedure not only avoids overfitting the data, but also enables us to discover novel features of motor adaptation in a powerful, exploratory fashion.

The best model predicted 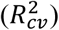 12.2%, 4.8%, and 18.6% of early, late, and aftereffect data, respectively. Our best model outperformed models built on randomly shuffled features (all p_perm_ < 0.001), underscoring its ability to predict motor behavior better than chance. However, the predictive capacity of a model based on individual differences was not extremely high; indeed, the variability in our dataset made it difficult to draw strong inferences on an individual level. As such, we focused on how learning functions differed on a cruder, categorical level. When the predictor was continuous (e.g., movement time), the learning functions were plotted based on a median split, binning participants into binary categories (e.g., high vs low movement times) (Table 2).

We start by focusing on features that predict early and late adaptation but are not predictive of the aftereffect (Figure 4a; a beta coefficient of 0 = not predictive). These features may be related to the participants’ efficiency in using strategic processes to counter the perturbation (e.g., re-aiming). First, the participant’s overall enjoyment rating of the experiment predicted early and late adaptation (Figure 4b). Whether adapting more causes greater enjoyment (i.e., more task success) or greater enjoyment elicits greater adaptation remains to be seen. Second, movement time predicted early and late adaptation, with faster movements times associated with greater adaptation (Figure 4c). Participants who moved faster may be those who were motivated to perform well (68). Alternatively, the strength of the error signal may weaken with movement time – an intriguing hypothesis that can be rigorously evaluated in the lab.

**Figure 4.**
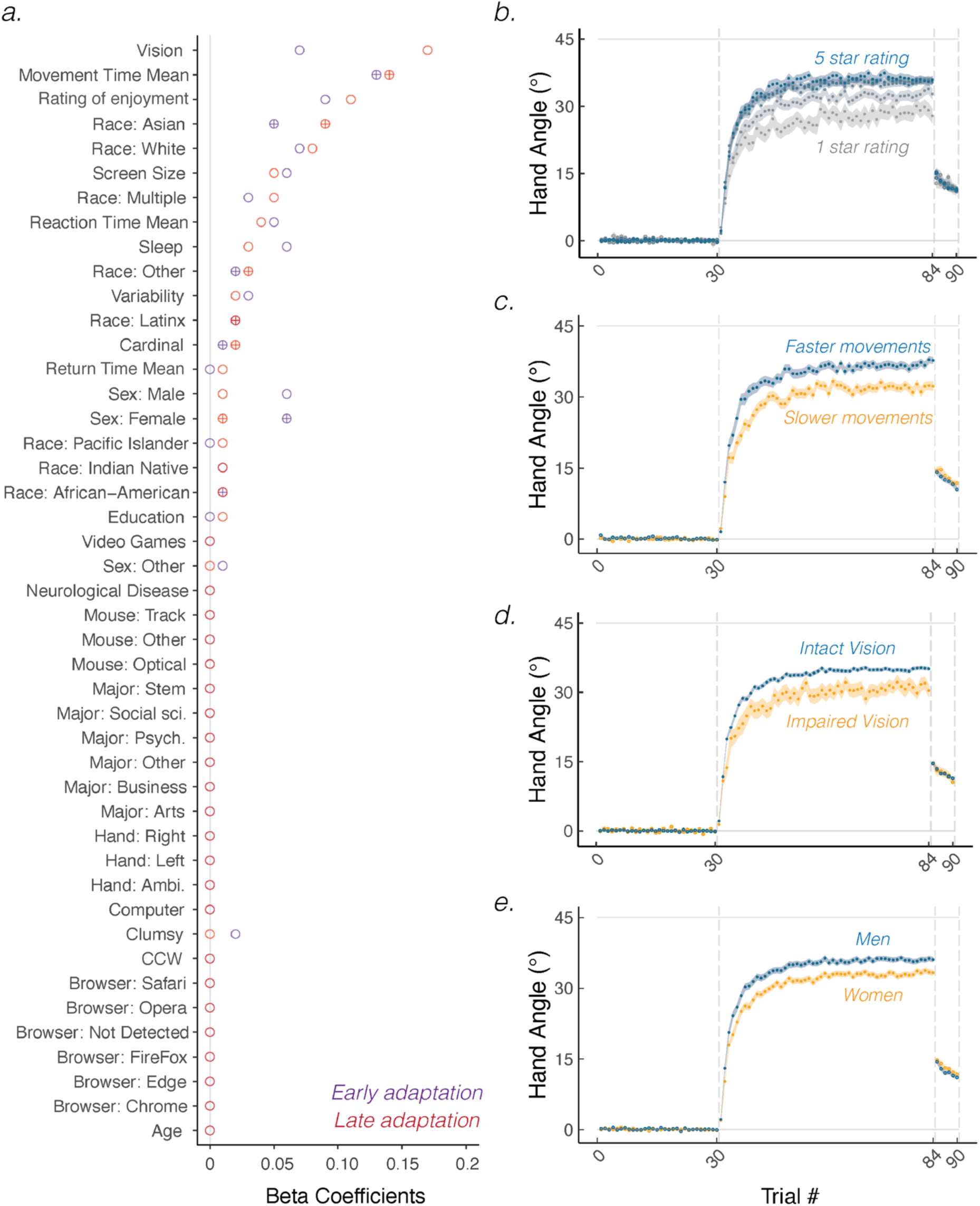
Predictors of early and late adaptation. **(a)** Beta coefficients indicate whether a feature positively (circle) or negatively (crosshairs) correlates with hand angle data. **(b-e)** Representative features that predict early and late adaptation: Rating of overall enjoyment, baseline movement time (median split), visual status, and sex. Shaded region denotes SEM.

Third, the participant’s sex (Figure 4e) predicted early and late adaptation. Compared to women, men were faster at counteracting the perturbation and reached a higher level of performance, consistent with what has been observed in in-person studies (69) (but see (67)). This result suggests that men are more likely to engage strategic processes to compensate for the visuomotor rotation. Fourth, participants who reported visual impairments adapted less than those who reported no visual impairments (Figure 4d). This finding suggests that successful strategic re-aiming may require high-fidelity visual input (also see Figure S1 for learning functions from other predictors).

### Identifying predictors of implicit recalibration

Next, we describe features that selectively predict variation in the size of the aftereffect (Figure 5a), our proxy of implicit recalibration. The effect of baseline motor variability on implicit recalibration has been a controversial topic (70,71). One perspective suggests that a more variable motor system may be sensitized to correct motor errors (72,73), and thus, would be associated with stronger implicit recalibration. Alternatively, it has been argued that large intrinsic noise reduces sensitivity to external perturbation by amplifying a “credit-assignment” problem (74–76). By this view, greater baseline variability would be associated with weaker implicit recalibration. Our results are consistent with the latter perspective: Higher baseline variability was associated with a lower asymptote (Figure 5b).

**Figure 5.**
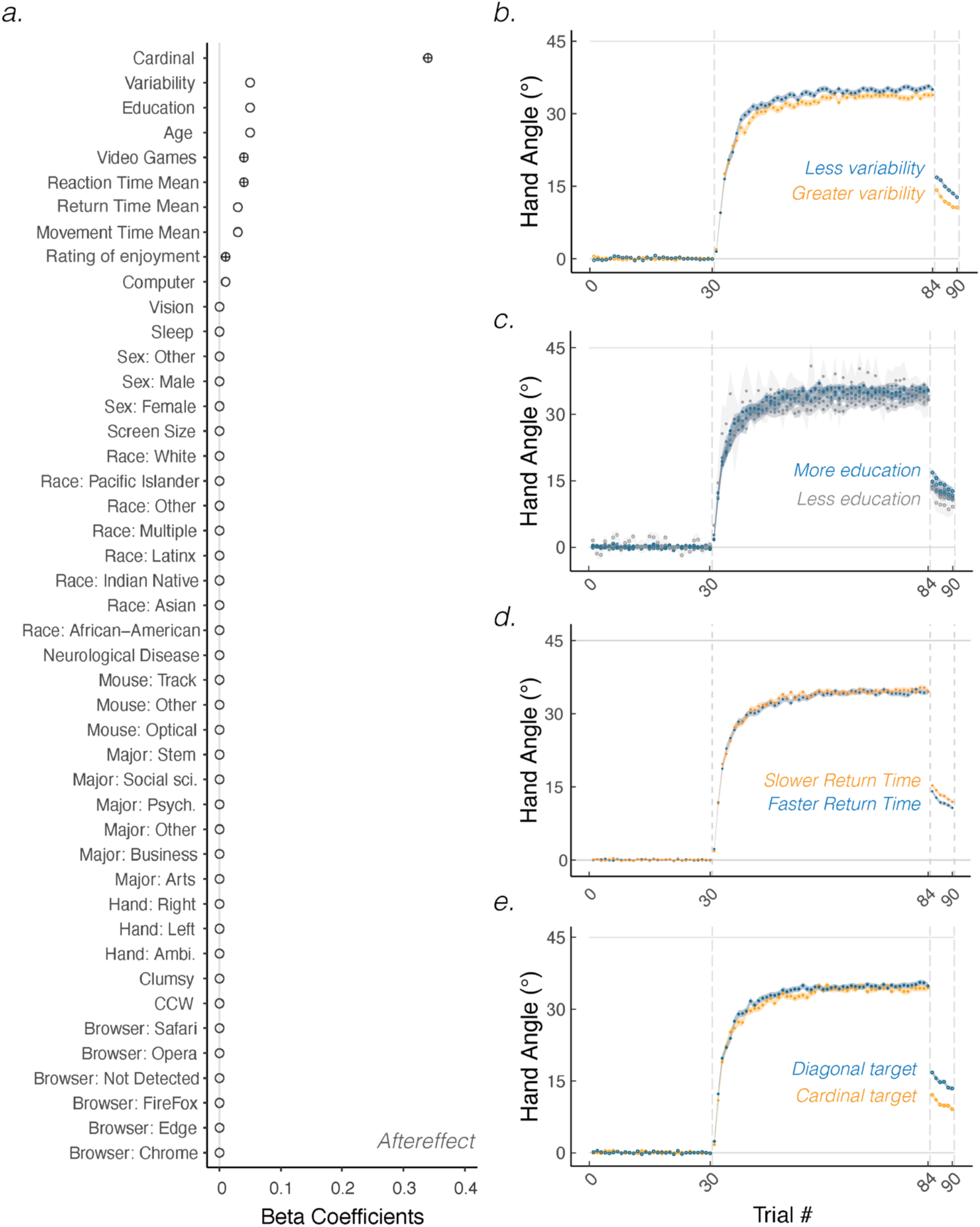
Predictors of motor aftereffect. **(a)** Beta coefficients indicate whether a feature positively (circle) or negatively (crosshairs) correlates with hand angle data. **(b-e)** Representative features that predict the size of the aftereffect: Baseline movement variability, level of education, baseline return time (median split), and target location. Shaded region denotes SEM.

Second, the participant’s level of education predicted the extent of the aftereffect (Figure 5c), with greater education associated with a larger aftereffect. This result may seem counterintuitive given that, *a priori*, we would expect education to be linked with cognitive variables such as strategy use. However, years of education is collinear with age (*r* = 0.53, *p* <0.001), a feature observed to modulate aftereffects (Figure 2b). As such, we suspect that those with more education may have been older, and thus exhibited slightly greater implicit recalibration (61,66,77–79).

Third, participant’s return time predicted the extent of aftereffects (Figure 5d), with longer baseline return times associated with a larger aftereffect. Participants who were slower to return their (hidden) cursor to the start position may be those who were less kinesthetically aware (i.e., less aware of where they had moved), a factor shown to correlate with the extent of implicit recalibration (46,80) (but see: (81,82)). Alternatively, longer return times may provide sufficient time for learning to consolidate (83), and as such, increase the extent of implicit recalibration.

Fourth, we observed an unexpected effect of target location on implicit recalibration. Participants who reached to diagonal target locations had an aftereffect that was almost twice as large as that observed for participants who reached towards cardinal targets. This result is especially noteworthy in that target location – being a variable of interest for motor control (i.e., different baseline movement biases) (84–86) – is often considered an nuisance variable for sensorimotor adaptation (with some exceptions, e.g., (87)). Future studies are required to evaluate the underlying mechanisms that give rise to these effects.

### Summary and Conclusions

Our data-driven web-based approach offers a powerful method to study sensorimotor learning outside the traditional laboratory (also see: (83,88–99)). First, we have shown that these data are reliable and valid, reproducing classic findings in the literature. For example, re-exposure to the same visuomotor rotation not only increased early and late adaptation (48,49,100–102) but also attenuated the aftereffect (45). Moreover, we observed the signature of contextual interference, in which blocked training (fixed target location) accelerated learning but weakened retention in comparison to training with multiple, interleaved targets.

Second, the large sample sizes possible in web-based studies offer a novel means to examine controversies in the sensorimotor learning literature. For example, we expect it may be difficult to detect the effects of age in lab-based studies with small sample sizes. Our results highlight a subtle, but striking, inverted-U effect of age, with late adaptation peaking between 20-50 years old and aftereffects peaking around 50-60 years old (also see: (67)). Future in-lab studies can home in on the mechanisms underlying these non-monotonic functions, asking how age-related cognitive decline may disrupt strategic re-aiming, and how age-related neural degeneration may impact implicit recalibration.

Third, our web-based approach allows us to tackle questions typically inaccessible to the lab and discover new predictors of sensorimotor adaptation. Leveraging this large dataset and a machine learning approach, we discovered that the participant’s sex, movement speed, and overall enjoyment of the experiment predicted the extent of strategic re-aiming. Age, level of education, return time, target location, and movement variability predicted the extent of implicit recalibration. Elucidating why these new, unappreciated features modulate different learning processes underlying sensorimotor adaptation will be an exciting area for future research.

There are notable limitations with this data-driven approach. Our predictive model only explained between 5% - 20% of the variance in the data. This may be because of the noisiness inherent in on-line data collection. One way to reduce this noise is to collect more data; doubling the number of trials would only add an additional 10 minutes to the current procedure. Our model is, of course, limited by our choice of measures. Future studies can build on the current model by assaying a wider range of features, including those we might expect to be predictive of motor performance (e.g., athleticism, musicality) as well as others we expect to be less predictive (e.g., geographic location, socio-economical background). These additions would take us closer to a more holistic model and understanding of sensorimotor learning.

## Methods

### Ethics statement

All participants gave written informed consent in accordance with policies approved by the UC Berkeley’s Institutional Review Board.

### Participants and Sessions

Participants were recruited between 2019 and 2022 on a citizen science website (TestMyBrain.org) that provides personalized performance feedback in exchange for study participation. A total of 2,289 experimental sessions were collected. For the visualization (Figure 1b-c), we excluded 168 sessions with erratic movements (i.e., the standard deviation of hand angle exceeded 25°, or more than 20% of outlier datapoints were removed; see Data Analysis) or systematic movements to the wrong direction (i.e., mean heading angle less than 0° or exceeded 75°), leaving 2,121 eligible sessions.

For the model-based analysis, we limited the data to participants who completed the one-target version of the task on their first session (n = 1,747). This criterion excluded 374 sessions (see Table 1) in which there may have been confounds (e.g., the two-target version impacted learning at all phases) and possible within-participant effects on behavior (e.g., savings or interference (45)). A summary of demographic and task features are provided in Table 2.

### Web-based sensorimotor adaptation task

All participants used their own laptop or desktop computer to access the TestMyBrain.org webpage that hosted the experiment (see a demo of the task at: https://multiclamp-c2.web.app/). Participants made reaching movements by moving the computer cursor with their mouse or trackpad. The size and position of stimuli were scaled based on each participant’s screen size. For ease of exposition, the stimuli parameters reported below are for a typical monitor size of 13” (1366 × 768 pixels), and the procedure reported below is for the one-target version of the task.

On each trial, the participants made a center-out planar movement from the center of the workspace to a peripheral target. The center position was indicated by a white annulus 0.5 cm in diameter, and the target location was indicated by a blue circle that was also 0.5 cm in diameter. The radial distance of the target from the start location was 6 cm. For each participant, the target always appeared at the same location on every trial. Each individual was randomly assigned a single target location selected from a set of eight possible locations (cardinal targets: 0°, 90°, 180°, 270°; diagonal targets: 45°, 135°, 225°, 315°).

To initiate each trial, the participant moved the cursor, represented by a white dot on their screen into the start location. During an introductory phase, feedback was only provided when the cursor was within 2 cm of the start circle. Once the participant maintained the cursor in the start position for 500 ms, the target appeared. The participant was instructed to reach to the target using the cursor. If the movement was not completed within 500 ms, the message “too slow” was displayed in red 20 pt. Times New Roman font at the center of the screen for 750 ms.

During the experimental phase, visual feedback could take one of the following forms: Veridical feedback, rotated feedback, and no feedback. During veridical feedback trials, the movement direction of the cursor was veridical with respect to the movement direction of the hand up to the target distance (6 cm). Once this distance was reached, the cursor position was frozen for 50 ms and then the cursor disappeared. During rotated feedback trials, the cursor moved at a 45° angular offset relative to the position of the hand up to the target distance (6 cm) before freezing for 50 ms. During no-feedback trials, the feedback cursor was extinguished as soon as the hand left the start circle and remained off for the entire movement. During the return phase after each movement, the veridical cursor was visible upon moving within 2 cm of the start circle.

Given the access demands for the testmybrain website, we were limited to only about 10 min of data collection. The study was designed to completed in under 10 minutes. As such, participants completed three blocks of trials (90 total trials): Baseline veridical-feedback block (30 trials), rotated-feedback block (54 trials), and a no-feedback block (6 trials). During the rotation block, the direction of rotation (i.e., clockwise, or counterclockwise) was counterbalanced across participants.

### Data analysis

The primary dependent variable was hand angle, defined as the angle of the hand relative to the target when the amplitude of the movement reached the target radius (6 cm). Positive hand angle values correspond to the direction opposite the rotated feedback (i.e., we flipped all hand angle values at targets where a counterclockwise rotation was provided).

The data were baseline subtracted. Baseline was defined as mean hand angle over all trials in the baseline block. Outlier trials were defined as trials in which the hand angle deviated by more than three standard deviations from a moving 5-trial window, or if the hand angle on a single trial was greater than 90° from the target. These trials were discarded since behavior on these trials likely reflects attentional lapses (average percent of trials removed: 1.6% ± 2.1%).

The degree of adaptation was quantified as the change in hand angle in the opposite direction of the rotation. We calculated hand angle during early adaptation, late adaption, and the aftereffect phase. Early adaptation was defined as the mean hand angle over the first 10 trials during the rotation block. Late adaptation was defined as the mean hand angle over the last 10 trials during the rotation block. Aftereffect was operationalized as the mean hand angle during the no-feedback block.

The hand angle data during early, late, aftereffect phases were entered into a group lasso regression as dependent variables (R function: cv.glmnet; (103)), and all the features in Table 2 were entered in as independent predictors. Categorical variables were assigned dummy variables (104); continuous variables were z-scored (105). We have opted to highlight results from group lasso regression since this procedure penalizes unimportant independent variables (i.e., sets them to zero). As such, the method is conservative in terms of identifying predictors. Group lasso also forces the model to keep or discard a pre-defined set of grouped variables (e.g., undergraduate major).

We used 10-fold cross validation on 80% of the sessions to select the model with the minimum mean cross-validation error. We fixed the best performing model’s beta coefficients and evaluated the degree to which this model predicted held out data (the remaining 20% of sessions). The absolute value of the beta-coefficient represents the importance of this feature in the model. We used the coefficient of determination 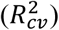 between the predicted and the actual held-out data as our key metric of model performance.

## Data availability statement

Data and code will be available upon publication at: https://github.com/itsalwaysnow/Identifying-predictors-of-sensorimotor-adaption-with-180-000-reaches.git.

---

**Figure S1.**
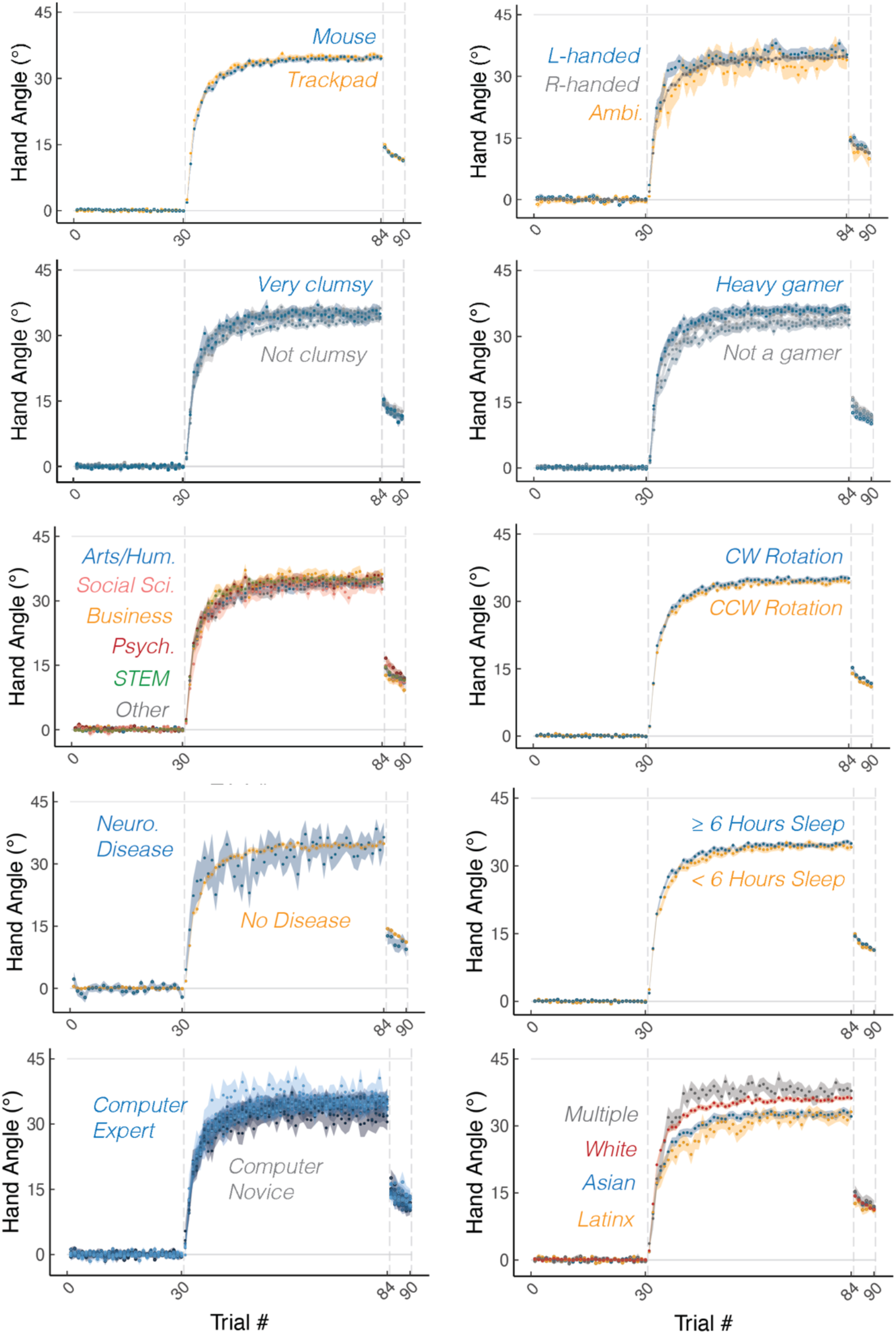
Other predictors of sensorimotor adaptation. Shaded region denotes SEM.

